# One-carbon metabolism enzyme Ahcy is a redox sensor that modulates gene expression to protect against light stress-induced retinal degeneration

**DOI:** 10.1101/2025.04.25.650714

**Authors:** Sarah C. Stanhope, Kratika Singhal, Nicolás M. Morato, Yunfei Feng, Gaoya Meng, Makayla N. Marlin, Claudia C. Kotanko, Madolyn M. Jarrett, Andrew D. Mesecar, Graham R. Cooks, Vikki M. Weake

## Abstract

One-carbon metabolism influences gene expression by providing methyl units for DNA, RNA, and histone methylation. Robust methylation requires rapid hydrolysis of the methylation by-product *S-*adenosylhomocysteine (SAH) by *S-*adenosylhomocysteinase (Ahcy). Here, we show Ahcy is a redox-sensitive enzyme that is inhibited by oxidation of a conserved cysteine, C195, *in vitro* and *in vivo*. Transient oxidation of Ahcy is neuroprotective in a *Drosophila* light stress model where it results in rapid gene expression changes and protects against retinal degeneration. Thus, redox sensing by the one-carbon metabolic enzyme Ahcy enables rapid changes in gene expression in response to changes in redox homeostasis.

## Introduction

The metabolic state of a cell is integrated with its transcriptional status via the epigenome because key metabolic intermediates also act as the donor molecules for deposition of epigenetic marks. In particular, one-carbon metabolism has a strong influence on the epigenome because histone, DNA, and RNA methylation require the universal methyl donor, *S*-adenosylmethionine (SAM), which is generated from methionine and folate (Serefidou et al. 2019). SAM availability impacts methylation of protein, DNA, and RNA based on the *K*_*m*_ values of each methyltransferase (Xiao et al. 2003; Patnaik et al. 2004; Obianyo and Thompson 2012; Chin et al. 2005; An et al. 2011). For example, histone H3 at lysine 4 trimethylation (H3K4me3) is sensitive to decreased SAM levels in embryonic stem cells (Shyh-Chang et al. 2013; Shiraki et al. 2014; Mentch et al. 2015). In addition, the methylation by-product *S*-adenosylhomocysteine (SAH) is a competitive and potent inhibitor of methyltransferase activity at relatively low concentrations (Catoni 1953; Glick et al. 1975). To prevent SAH inhibition and enable high methyltransferase activity, SAH is rapidly hydrolyzed by *S*-Adenosylhomocysteinase (Ahcy; also known as *S*-adenosylhomocysteine hydrolase, SAHH), to homocysteine and adenosine (Palmer and Abeles 1979). Ahcy is the sole enzyme that catalyzes SAH hydrolysis and is one of the most conserved enzymes among higher eukaryotes (Vizán et al. 2021). The homocysteine generated by SAH hydrolysis is either remethylated as part of the methionine salvage cycle or is converted to glutathione via the transsulfuration pathway (Ducker and Rabinowitz 2017). In addition to its cytoplasmic activity, Ahcy is found in the nucleus, where it associates with chromatin on the promoters of actively transcribed genes (Aranda et al. 2019). Moreover, Ahcy is necessary in mouse embryonic fibroblasts for both the cyclic deposition of H3K4me3 and circadian transcription of many rhythmic genes (Greco et al. 2020). Thus, Ahcy acts at a critical regulatory node in one-carbon metabolism that controls flux to support redox homeostasis and promotes the methylation reactions that regulate gene expression (Ducker and Rabinowitz 2017).

Using redox proteomics, we previously identified a potential redox-sensitive cysteine residue in Ahcy that exhibited changes in cysteine availability, indicative of its oxidation, in *Drosophila* eyes exposed to light stress (Stanhope et al. 2023). In *Drosophila* eyes, exposure to prolonged blue light alters redox homeostasis, decreasing the ratio of reduced to oxidized glutathione, and increasing hydrogen peroxide (H_2_O_2_) levels and lipid peroxidation, so resulting in eventual photoreceptor death and retinal degeneration (Escobedo et al. 2022; Chen et al. 2017; Stanhope et al. 2023; Hall et al. 2018). Prior to degeneration, photoreceptors exposed to light stress undergo a neuroprotective gene expression program (Hall et al. 2018); however, the gene regulatory mechanisms responsible for the rapid changes in gene expression remain unknown. Since the oxidized cysteine we identified in Ahcy is necessary for full activity of the human enzyme *in vitro* (Yuan et al. 1996), we hypothesize that Ahcy acts as a redox-sensitive enzyme that can rapidly alter gene expression programs in response to changes in redox homeostasis, potentially by altering histone methylation. Here, we show that transient inhibition of Ahcy enzymatic activity under oxidizing conditions, specifically light stress, leads to gene expression changes that protect against neurodegeneration. However, these gene expression changes do not correlate with Ahcy-dependent changes in H3K4me3 levels. Thus, methylated histones may not be the relevant target for Ahcy’s impact on gene expression. Since Ahcy is also oxidized in H_2_O_2_-treated *Drosophila* cultured S2 cells (Stanhope et al. 2023) and mouse kidney epithelial cells (Van Der Reest et al. 2018), this redox-sensing role of Ahcy may be broadly relevant to multiple tissues across species. Our data support a model in which changes in redox homeostasis can rapidly alter cellular gene expression programs by altering activity of this key regulatory enzyme in one-carbon metabolism.

## Results and Discussion

### Oxidation of Ahcy at C195 correlates with increased SAH

We previously observed decreased iodoacetamide-TMT labeling of C195 in Ahcy in *Drosophila* eyes exposed to light stress by using redox proteomics, indicative of cysteine oxidation (Stanhope et al. 2023). Two independent peptides containing C195 significantly decreased in abundance upon light stress without changes in abundance of total Ahcy protein or its putative regulator AhcyL1 (Fig. 1A) (Stanhope et al. 2023). Ahcy is the sole enzyme that hydrolyzes SAH into adenosine and homocysteine (Turner et al. 1998), and is necessary to prevent SAH-inhibition of methyltransferases (Hildesheim et al. 1973) (Fig. 1B). Ahcy is highly conserved (Kusakabe et al. 2015) and C195 is a critical residue for catalytic activity of the human enzyme (Yuan et al. 1996). If oxidation of Ahcy at C195 inhibits its activity, we would expect to observe increased SAH levels upon light stress. When we measured SAM and SAH in blue light-treated flies, we observed a significant increase in SAH without any corresponding change in SAM (Fig. 1C). We observed similar increases in SAH in S2 cells treated with H_2_O_2_, where Ahcy C195 oxidation is also observed (Fig. 1D) (Stanhope et al. 2023). To examine Ahcy oxidation in flies, we labeled available cysteines in wild-type (WT) and C195S Ahcy using MM(PEG)_24_, which increases the molecular weight (MW) by 1.24 kDa for each conjugated cysteine residue (Pant et al. 2021). We observe a small decrease in the MW of C195S mutant relative to WT Ahcy, consistent with loss of one cysteine labeled residue (Fig. 1E). Moreover, there is a small fraction of WT, but not C195S mutant, Ahcy that exhibits a decrease in MW consistent with loss of availability of two cysteine residues (Fig. 1E). The absence of this lower MW band in the C195S mutant indicates that the oxidation of the second cysteine depends on the presence of C195. These data show that under normal light conditions, a small fraction of Ahcy is oxidized at C195, and that this oxidation event may involve formation of a disulfide bond with a second nearby cysteine. Thus, oxidation of Ahcy at C195 correlates with increased SAH levels *in vitro* and *in vivo*.

**Figure 1.**
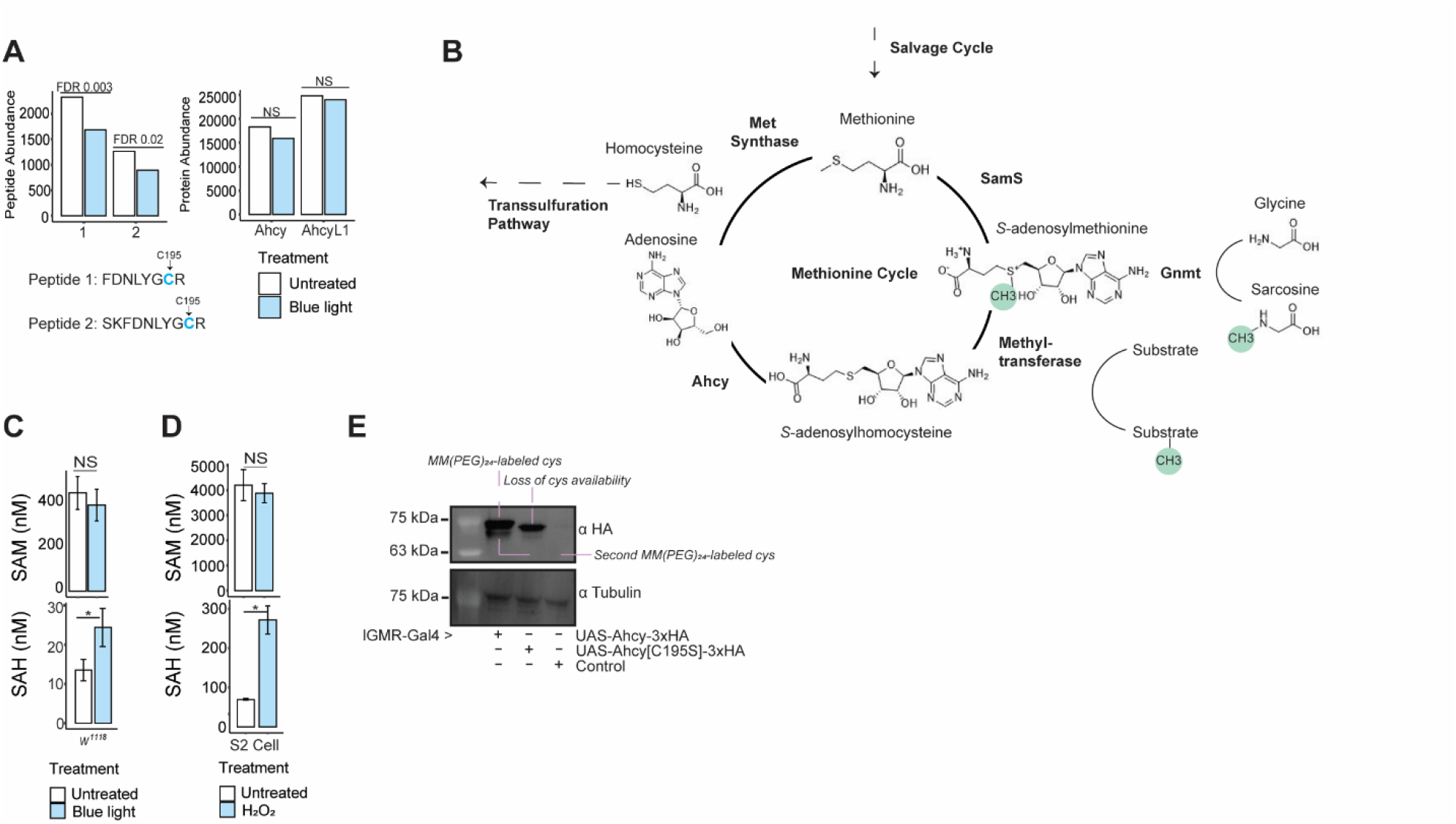
Ahcy oxidation correlates with increased SAH in cells and flies. *(A)* Bar graph of Ahcy iodo-TMT labeled peptide abundance (left) and Ahcy and AhcyL1 protein abundance (right) from *Drosophila* eyes exposed to blue light relative to untreated controls; Data for Ahcy graphed from published redox proteomic study (Stanhope et al. 2023). Iodo-TMT labeled peptides 1 and 2 containing cysteine 195, indicated in *blue*, showed significantly decreased peptide abundance upon blue light exposure indicative of decreased cysteine availability. False Discovery Rate (FDR) < 0.05 determined from PolySTest. *(B)* Schematic of selected enzymes and reactions involved in one-carbon metabolism; methyl groups indicated in *green. (C, D)* Bar graph displaying SAM and SAH levels in fly heads exposed to blue light relative to untreated controls (panel C) and S2 cells treated with 20 mM H_2_O_2_ (panel D). Graph depicts mean ± standard deviation (S.D.) (n = 3). *p-value* (* < 0.05) Student’s t test. (E) Western blot of MM(PEG)_24_ labeled protein extracts from fly heads overexpressing WT Ahcy or C195S mutant Ahcy.

### Ahcy is not regulated by direct AhcyL1 binding

Our data support the hypothesis that oxidation of Ahcy at C195 inhibits its activity *in vivo*. However, an alternative model has been proposed in which formation of a heterotetramer between Ahcy and its non-catalytic paralog AhcyL1 inhibits its activity (Devogelaere et al. 2008). Interactions between Ahcy and AhcyL1 have been reported in cultured *Drosophila* S2 cells and mammalian cells using co-immunoprecipitation or bimolecular fluorescence complementation (BiFC), respectively (Parkhitko et al. 2016; Grbeša et al. 2017). In our data, the increased SAH levels that correlated with Ahcy C195 oxidation were not accompanied by increases in AhcyL1 protein abundance (Fig. 1A) (Stanhope et al. 2023). We also did not observe any increase in abundance of a second Ahcy paralog, AhcyL2. Thus, we revisited the question of whether Ahcy was in fact inhibited by AhcyL1 binding. We co-immunoprecipitated epitope-tagged Ahcy, AhcyL1, and AhcyL2 in *Drosophila* S2 cells, but were unable to detect any interaction between Ahcy and either paralog (Fig. 2A). Despite this, both Ahcy and AhcyL1 could interact robustly with themselves, consistent with homotetramer formation (Fig. 1A & Supp. Fig. 1). We observed similar results by yeast two-hybrid where we did not detect any interactions between Ahcy, AhcyL1, and AhcyL2, even though Ahcy and AhcyL1 could interact with themselves (Fig. 2B). Thus, in contrast with the previous reports of an interaction between Ahcy and AhcyL1 (Parkhitko et al. 2016; Grbeša et al. 2017), our data do not support a model in which Ahcy is negatively regulated by formation of a heterotetramer with AhcyL1 and raise doubts as to whether these proteins can in fact interact with each other.

**Figure 2.**
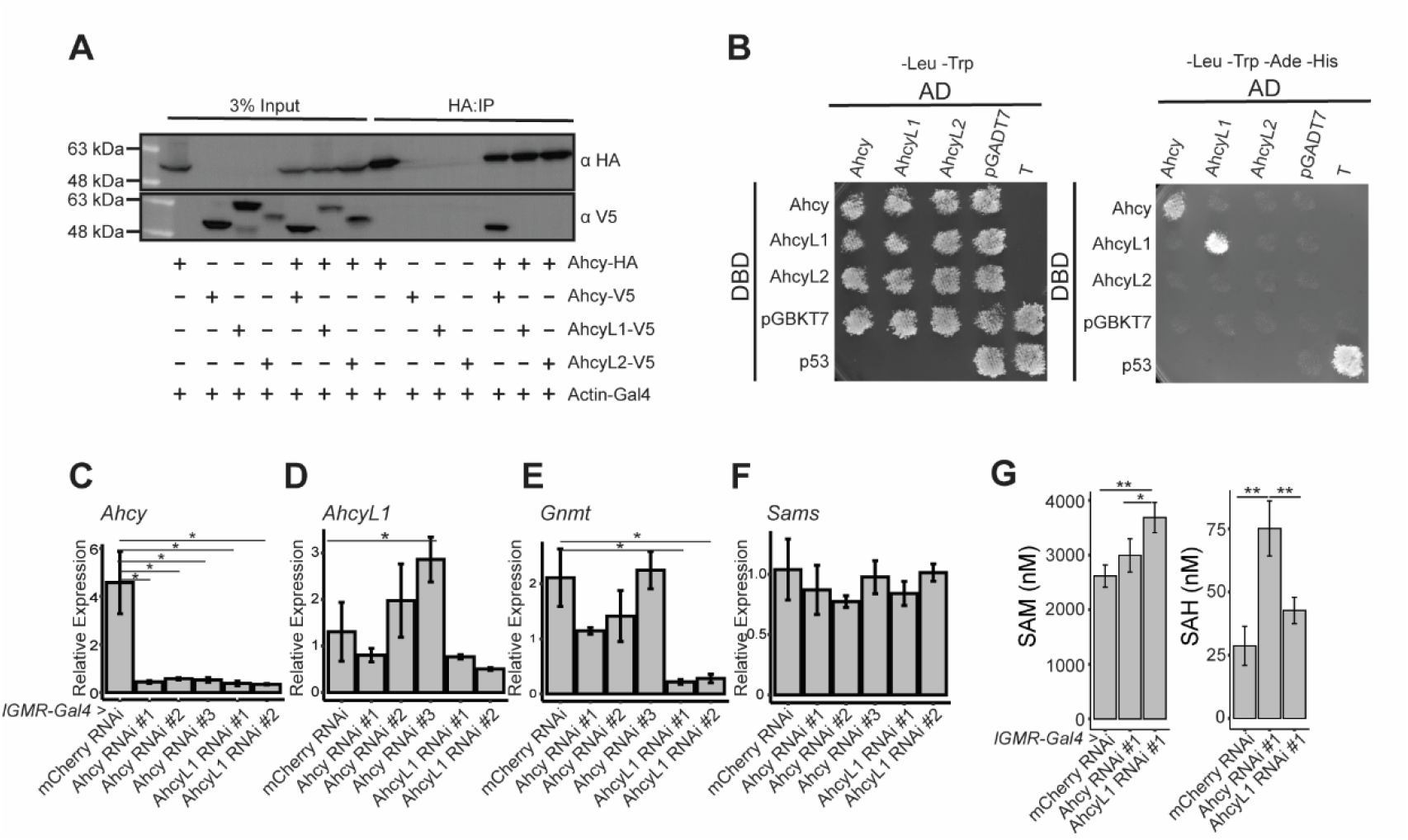
Ahcy is not regulated by binding of AhcyL1. *(A)* Co-immunoprecipitation of overexpressed epitope-tagged Ahcy, AhcyL1, and AhcyL1 in S2 cells. *(B)* Yeast two-hybrid analysis of Ahcy, AhcyL1, and AhcyL2 interactions. Growth on -leu -trp -ade -his media indicates positive interaction, with p53/T as positive control. AD, Gal4 activating domain; DBD, Gal4 DNA binding domain. *(C-F)* Bar graphs of relative expression of the indicated genes as determined by RT-qPCR from eye-specific *lGMR-Gal4*>RNAi of the specified lines relative to the control, mCherry RNAi. Relative expression was normalized to *Rpl32* and *Eif1a*. Graphs depict mean ± S.D. (n = 3). *p-value* (* < 0.05) Student’s t-test. *(G)* Bar graph displaying SAM and SAH levels in fly heads from eye-specific *lGMR-Gal4*>RNAi for the indicated lines. Graph depicts mean ± S.D. (n = 3). *p-value* (* < 0.05, ** < 0.01) ANOVA with Tukey’s posthoc.

Based on our failure to replicate the Ahcy-AhcyL1 interaction, we wondered about the interpretation of studies based on this regulatory model that have shown its importance for lifespan extension. Previous studies used RNAi in *Drosophila* to show that tissue-specific downregulation of AhcyL1/L2 extends lifespan by modulating Ahcy activity (Parkhitko et al. 2016). We examined the same two AhcyL1 RNAi lines used in that study to knockdown AhcyL1 in flies, and found that both lines significantly decrease Ahcy transcript level with only modest effects on AhcyL1 (Fig. 2C & D). In contrast, three independent Ahcy RNAi lines all decrease Ahcy expression without any corresponding changes in AhcyL1, or in other one-carbon metabolism enzymes such as Glycine-N-methyltransferase (Gnmt) and S-adenosylmethionine Synthetase (Sams) (Fig. 2E & F). Whereas Ahcy knockdown in eyes increases SAH levels but does not affect SAM, AhcyL1 knockdown slightly increases SAM but not SAH (Fig. 2G). Even though AhcyL1 knockdown decreased Ahcy transcript levels, it did not increase SAH. We attribute the failure of AhcyL1 knockdown to increase SAH levels to the concomitant decrease in both Ahcy and Gnmt expression (Fig. 2C & E). Gnmt methylates glycine to form sarcosine, thereby regulating SAM and SAH levels (Fig. 1B) and has a role in lifespan extension in flies (Obata and Miura 2015). We conclude that Ahcy is not regulated by AhcyL1 binding, leading us to pursue our hypothesis that oxidation of Ahcy at C195 negatively regulates its enzymatic activity.

### Oxidation of Drosophila Ahcy inhibits enzyme activity in vitro

We purified recombinant WT and C195S mutant *Drosophila* Ahcy and compared their hydrolytic activity *in vitro* using a **<u>H</u>**igh-**<u>T</u>**hroughput **<u>D</u>**esorption **<u>E</u>**lectro**<u>s</u>**pray **I**onization **<u>M</u>**ass **<u>S</u>**pectrometry (HT-DESI-MS) to directly quantify adenosine production (Supp. Fig. 2A) from the bioassay mixture (Morato et al. 2020, 2021). We observed strong substrate inhibition (i.e., SAH) in initial reaction velocities for both *Drosophila* WT and C195S mutant (Fig. 3A & B), which has been reported by previous studies (Walker and Duerre 1975; Kailing et al. 2017). The C195S mutation substantially decreases enzymatic activity as reflected by the order of magnitude reduction in the turnover number (*k*_*cat*_) of the mutant (0.028 ± 0.003 s^-1^) compared to that of the WT (0.34 ± 0.07 s^-1^). Additionally, the C195S showed a 4-fold difference in its catalytic efficiency (*k*_*cat*_/*K*_*m*_) which decreases from 8 ± 2 nM s^-1^ in the WT Ahcy to 2.1 ± 0.4 nM s^-1^ (Fig. 3A & B). A similar decrease in activity has been reported for human C195S Ahcy (Yuan et al. 1996). Interestingly, although C195 is important for *Drosophila* and human Ahcy activity, and is conserved among nearly all Ahcy homologs, *Caenorhabditis elegans* Ahcy has an isoleucine rather than a cysteine at this position (Fig. 3C). Since Ahcy plays a critical role in SAH hydrolysis in all organisms, we wondered whether *C. elegans* Ahcy would exhibit comparable enzymatic activity despite the lack of this highly conserved C195 residue. To test this, we purified recombinant *C. elegans* Ahcy and examined its enzymatic activity *in vitro* via HT-DESI-MS (Fig. 3B). Interestingly, the worm enzyme showed a catalytic efficiency of 5 ± 1 nM s^-1^, similar to that of WT *Drosophila* Ahcy, demonstrating that it is enzymatically active *in vitro* despite the lack of this conserved C195. These data indicate that C195 could have a role in regulating Ahcy enzymatic activity in flies and humans, but not in worms.

**Figure 3.**
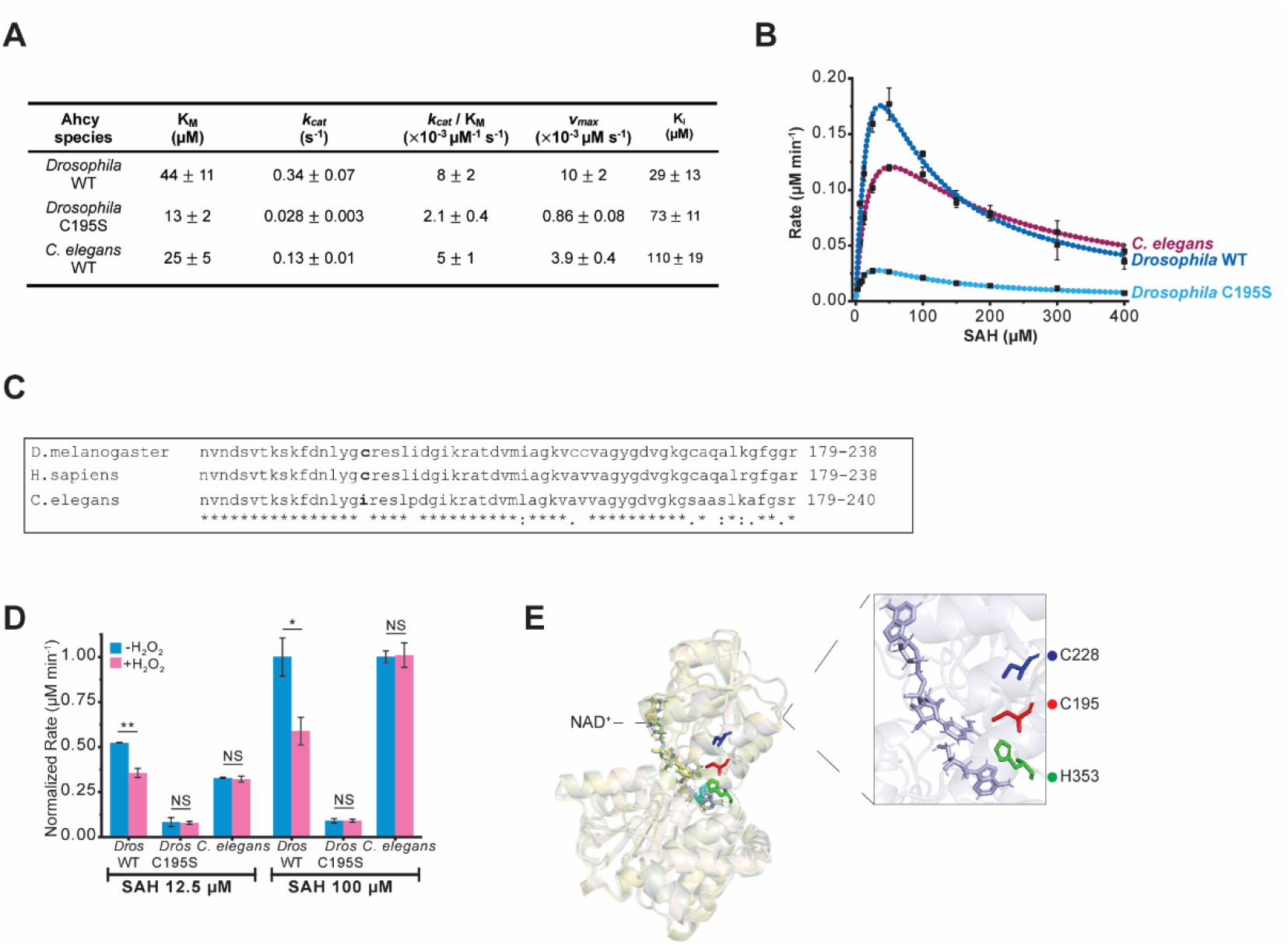
*Drosophila* Ahcy enzymatic activity is inhibited by oxidation of C195. *(A)* Table of kinetic characterization of *Drosophila* wild-type (WT) Ahcy, *Drosophila* C195S mutant Ahcy, and *C. elegans* Ahcy as determined by DESI-MS analysis. *(B)* Graph of kinetic characterization of indicated enzymes plotted using substrate inhibition model. Graph depicts mean ± S.D. (n = 3). *(C)* Alignment of selected region of Ahcy from the indicated species using Clustal Omega with conserved cysteine 195 residue bolded. An asterisk (*) indicates positions which have a single, fully conserved residue, a colon (:) indicates a conversation between groups of strongly similar properties, and period (.) indicates a conversation between groups of weakly similar properties. *(D)* Bar graphs displaying normalized rate of indicated enzymes with and without pre-incubation of H_2_O_2_ at SAH concentrations of 12.5 μM and 100 μM. All *Drosophila* enzymes were normalized to the untreated *Drosophila* WT Ahcy. *C. elegans* samples were normalized to the untreated *C. elegans* WT Ahcy. Graph depicts mean ± S.D. (n = 3). *p-value* (* < 0.05, ** < 0.01) Student’s t-test. *(E)* X-ray crystal structure of superimposed *Drosophila* WT Ahcy (*left)* and a single WT Ahcy subunit (*right)* highlighting C195, C228, H353, D190, and NAD^+^ redox cofactor at 2.7 Å.

We next asked if oxidation of *Drosophila* Ahcy inhibited its enzymatic activity *in vitro*. Pre-incubation of WT *Drosophila* Ahcy with H_2_O_2_ significantly decreased reaction velocity to about 50% of untreated controls at two different SAH concentrations (Fig. 3D). In contrast, C195S mutant showed no difference between H_2_O_2_-treatment and control, indicating that oxidation of this specific cysteine residue is responsible for the decrease in enzymatic activity. If oxidation of C195 in Ahcy inhibits its activity, then we predicted that the worm Ahcy homolog would be redox-insensitive. Indeed, *C. elegans* Ahcy showed no changes in activity between H_2_O_2_-treatment and control. Thus, we confirmed that oxidation of Ahcy at C195 inhibits its enzymatic activity.

To understand how C195 oxidation inhibits Ahcy activity, WT *Drosophila* Ahcy was co-crystallized with NAD^+^ and its X-ray structure determined to 2.7 Å (Fig. 3E; Supp Table 3). The resulting asymmetric unit contained all four monomers of the Ahcy biological tetramer, and each monomer was bound with NAD^+^ (Supp Fig. 2B). Adenosine was also present in three of the subunits although it was not included in the crystallization solution, indicating that endogenous adenosine from *E*.*coli* remained tightly bound to the enzyme during purification. The side chain of C195 hovers directly over the hydrogen atom of the C4 atom in the NAD^+^ nicotinamide ring with an S to C4 distance of 3.6 Å (Fig. 3E). The C195 thiol is within hydrogen bonding distance of the E2-nitrogen atom H353 (3.4Å) that also forms a hydrogen bond with the D190 sidechain carboxylate group. D190 is postulated to be involved in proton abstraction from K185 which in turn abstracts a proton from the 3’-OH on the adenosine ribose ring during hydride transfer (Yamada et al. 2005).

Although a formal role for C195 in the Ahcy mechanism has yet to be defined, it is clearly involved in catalysis given the 12-fold reduction in *k*_cat_ and the 4-fold reduction in *k*_cat_/K_m_ observed with C195S (Fig. 3B). One likely role for C195 is to properly position residues H353, D190, and K185 to conduct proton abstraction during hydride transfer from C3 to NAD^+^ forming NADH. Even slight structural alterations generated by C195S could generate a significant reduction in catalysis (Mesecar et al. 1997). Another potential role for C195 would be to assist in the local redox potential surrounding NAD^+^/NADH during catalysis. Upon SAH binding, Ahcy utilizes NAD^+^ to oxidize the 3’OH group of the ribose sugar, generating NADH (Palmer and Abeles 1979). SAH is subsequently cleaved to form homocysteine and 3’keto-adenosine, which is reduced to adenosine using NADH. NADH also provides the 3’OH reduction potential necessary for the release of adenosine and homocysteine. Previous reports have suggested that C195 is necessary for the hydride transfer to regenerate NAD^+^ from NADH (Yuan et al. 1996), and our structure and kinetic studies are consistent with this model.

Lastly, when Ahcy is bound with NAD^+^, C195 is completely protected from solvent (i.e., H_2_O_2_) suggesting that any oxidation of C195 would have to occur when NAD is not bound. Alternatively, if the nicotinamide ring of bound NAD has some dynamic motion, this could allow penetration by an oxidant. With either mechanism, oxidation of C195 would be expected to significantly disrupt enzymatic catalysis perhaps even more so than the C195S substitution. These data support our hypothesis that in *Drosophila* and mammals where this C195 residue is conserved, Ahcy is a redox-regulated enzyme that is inhibited by reversible oxidation of C195.

### Ahcy is necessary for the light stress gene expression response

In addition to its cytoplasmic roles, Ahcy localizes to chromatin where it is necessary for gene expression, which has been attributed in part to promotion of H3K4me3 (Aranda et al. 2019; Greco et al. 2020). Flies exposed to blue light stress exhibit gene expression changes in photoreceptor neurons as part of a neuroprotective response, but the mechanisms involved in these transcriptional changes are unknown (Escobedo et al. 2022; Hall et al. 2018). If light stress results in oxidation and inhibition of Ahcy, then the subsequent build-up in SAH levels could decrease activity of methyltransferases, potentially including histone methyltransferases, thereby leading to global changes in gene expression. To test if Ahcy was required for light stress-dependent changes in gene expression, we profiled the photoreceptor nuclear transcriptome of Ahcy RNAi flies exposed to light stress relative to RNAi control (mCherry RNAi) (Fig. 4A). Principal Component Analysis revealed that the light stress control RNAi groups resembled Ahcy RNAi untreated samples (Fig. 4B), consistent with inhibition of Ahcy upon light stress in control RNAi flies. Moreover, while ~500 genes were differentially expressed between light stress and untreated samples in the control RNAi flies, fewer than 100 were differentially expressed upon light stress in the Ahcy RNAi flies (Fig. 4C & supp Table 4). In control RNAi flies, light stress-induced genes include factors involved in neuronal adaptation and chronic stress response such as neuropeptides (*snpf, npf, nplp1)* and genes needed for calcium signaling including *cam, canA-14f*, and *canB* (Fig. 4C). Prolonged calcium influx during light stress leads to retinal degeneration, which can be suppressed by mutations in the Trp calcium channel (Chen et al. 2017), but calcium influx can also alter gene expression. Moreover, the genes that were downregulated upon light stress in control RNAi flies include several factors that regulate phototransduction such as *inaF* as well as stress response genes including the apoptotic factor *dcp-1* (Fig. 4C). Mutations in *inaF* decrease abundance of the Trp calcium channel (Li et al. 1999); thus, decreased *inaF* expression could provide a mechanism to downregulate the light stress-dependent calcium influx to protect against calcium excitotoxicity. Ahcy knockdown suppresses these light stress-regulated gene expression changes, largely because those light stress-regulated genes already show differential expression between the untreated control RNAi and Ahcy RNAi flies (Fig. 4D & Supp. Fig. 3). For instance, the light stress-induced genes were already expressed at higher levels in untreated Ahcy RNAi flies versus control RNAi (Fig. 4D). Similarly, the light stress-repressed genes were already at lower levels in untreated Ahcy RNAi flies versus control RNAi. These data are consistent with a model in which light stress inhibits Ahcy via oxidation, leading to changes in gene expression that enable photoreceptors to adapt to prolonged phototransduction.

**Figure 4.**
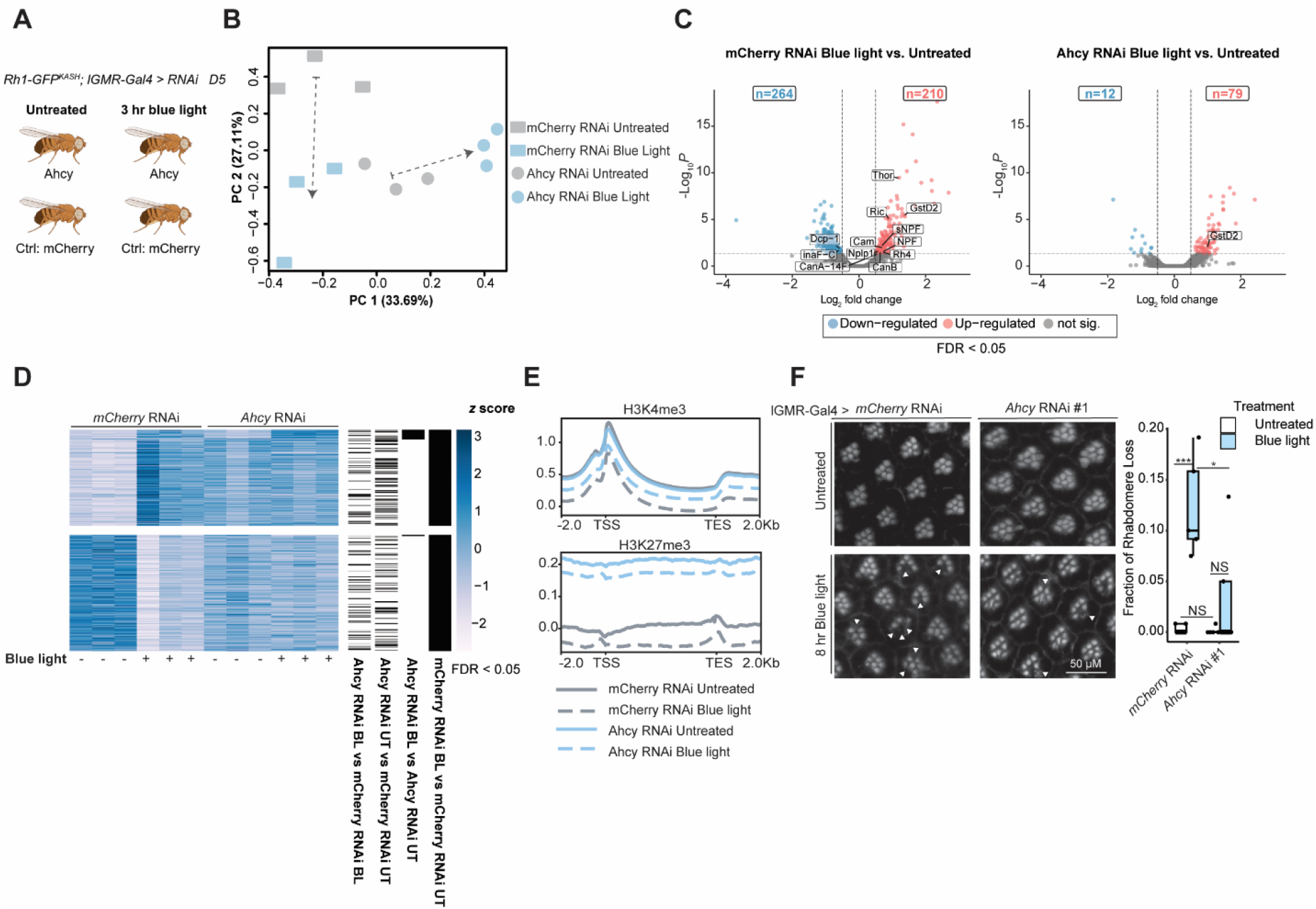
Ahcy is necessary for the light stress gene expression response and neuroprotection. *(A)* Schematic of RNAseq and CUT&RUN experimental design. Photoreceptor nuclei were labeled using *Rh1-GFP*^*KASH*^ in flies expressing eye-specific *lGMR-Gal4*>RNAi, that were exposed to 3h blue light versus white light control. *(B)* Principal Component Analysis (PCA) of RNAseq samples based on variance-stabilized gene expression levels (n = 3). *(C)* Volcano plots displaying significantly up- and down-regulated genes in either *mCherry* RNAi flies or *Ahcy* RNAi flies exposed to blue light relative to untreated controls. FDR < 0.05, log_2_(fold-change) ≥ 1 or ≤ −1). *(D)* Heatmap showing genes that were significantly differentially expressed upon blue light in *mCherry* RNAi control flies. Colors indicate *z* score of relative expression in biological replicates for each genotype and treatment. All genes shown are differentially expressed genes (DEGs) from *mCherry* RNAi flies exposed to blue light relative to untreated controls. Other categories shown include *Ahcy* RNAi flies exposed to blue light relative to untreated controls, *Ahcy* RNAi untreated flies relative to *mCherry* RNAi untreated flies, and *Ahcy* RNAi flies exposed to blue light relative to *mCherry* RNAi flies exposed to blue light. *(E)* Gene metaplots representing mean normalized counts for CUT&RUN of indicated histone methyl marks in each genotype and treatment (n = 3). Gene bodies are shown from TSS (transcription start site) to TES (transcription end site) for all actively expressed unique transcription start sites. *(F)* Representative images of phalloidin stained blue light exposed or untreated eyes from indicated genotypes. Box plots showing quantification (displayed as fraction of rhabdomere loss) of R1-R6 rhabdomeres from indicated genotypes. Arrow heads point to ommatidia in the eye with degenerated rhabdomeres. Box plots with overlaid points were generated using ggplot2; lower and upper hinges correspond to the first and third quartiles, and the whiskers extend to the smallest or largest values no more than 1.5x inter-quartile range (IQR) from each hinge. Graph depicts mean ± S.D. (n = 5). *p-value* (* < 0.05, ** < 0.01, *** < 0.001) ANOVA with Tukey’s posthoc.

Ahcy has been implicated previously in both gene activation and repression by promoting methylation of different histone lysine residues. For instance, while Ahcy-dependent enhancement of H3K4me3 is associated with transcription activation (Aranda et al. 2019; Greco et al. 2020), Ahcy inhibition in mouse embryonic stem cells decreases levels of the repressive H3K27me3 mark (Jiang et al. 2024). However, Ahcy appears to have context-dependent roles in histone methylation because Ahcy knockout or inhibition via 3-deazaneplanocin A (DZnep) in mouse embryonic fibroblasts did not alter H3K27me3 (Greco et al. 2020). DZnep is a specific inhibitor of Ahcy activity that is often referred to as an epigenetic inhibitor due to its impact on histone methylation (Miranda et al. 2009). We examined H3K4me3 and H3K27me3 levels in photoreceptors from Ahcy RNAi flies exposed to blue light relative to RNAi control using CUT&RUN. H3K4me3 levels decreased upon blue light exposure in both Ahcy RNAi and mCherry RNAi control flies, and did not substantially differ between the untreated mCherry RNAi control and Ahcy RNAi flies (Fig. 4E). Interestingly, H3K27me3 levels actually increased in Ahcy RNAi untreated and blue light exposed flies relative to the mCherry RNAi flies, with only slight decreases in H3K27me3 in both genotypes upon blue light treatment (Fig. 4E). Thus, neither H3K4me3 nor H3K27me3 correlate with the suppression of light stress-dependent gene expression changes by Ahcy knockdown, indicating that these are unlikely to be the relevant targets that mediate transcriptional changes dependent on Ahcy oxidation. We note that previous studies have shown that H_2_O_2_ treatment decreases H3K4me3 levels in mammalian HeLa cells, which was attributed to redox sensing by the methyltransferase Set1 (Bazopoulou et al. 2019); however, we did not identify Set1 as a potentially oxidized protein in *Drosophila* cells or eyes in our redox proteomic analysis (Stanhope et al. 2023). Our findings suggest that redox sensing by Ahcy is required for rapid changes in transcription caused by light stress in photoreceptors, but this is unlikely mediated by alterations in the two histone methylation marks previously shown to be dependent on Ahcy activity.

### Ahcy knockdown protects against light stress-induced retinal degeneration

Since Ahcy oxidation under light stress leads to rapid gene expression changes enabling neuronal adaptation to this prolonged activation state, we hypothesized that the inhibition of Ahcy activity under light stress was neuroprotective. To test this, we exposed Ahcy RNAi and control RNAi flies to prolonged blue light and assessed retinal degeneration by staining retinas with phalloidin, which marks the actin-rich rhabdomeres. As expected from previous studies (Chen et al. 2017), there was a significant decrease in the number of rhabdomeres, indicative of photoreceptor loss, in control RNAi flies exposed to blue light versus untreated flies (Fig. 4F). Strikingly, Ahcy RNAi flies exposed to blue light exhibited much lower levels of rhabdomere loss when compared both with untreated Ahcy RNAi flies and with mCherry RNAi flies exposed to blue light (Fig. 4F). Thus, knockdown of Ahcy protected eyes against light stress-induced retinal degeneration, indicating that the transient oxidation and inhibition of Ahcy activity upon light stress is neuroprotective.

Overall, these data demonstrate that Ahcy is a redox sensor that can rapidly alter gene expression programs in response to changes in redox homeostasis, thereby directly connecting metabolism and transcription. In the *Drosophila* eye, this transient inhibition of Ahcy activity is neuroprotective against light stress. Although several specific methylated histone marks have previously been implicated in Ahcy-dependent gene expression changes, our data indicate that Ahcy may instead promote methylation of an as-of-yet uncharacterized target critical for transcription regulation. Since the redox-sensitive cysteine in Ahcy is conserved in mammals, redox-sensing by this enzyme involved in one-carbon metabolism may be a widespread mechanism to rapidly modulate gene expression in response to changes in redox homeostasis across tissues.

## Materials and Methods

### *Drosophila* Stocks and Culture

*Drosophila* were maintained at 25°C under 12-h:12-h light:dark conditions where ZT0 (ZT; zeitgeber time) indicates the beginning of the light phase. Male flies were exposed to 3 h blue light or white light control, as in (Escobedo et al. 2022), beginning at ZT9 at 5 days post-eclosion, and collected at ZT12. Stocks for light stress were crossed into the *cn bw* background to deplete eye pigments. *UAS-Ahcy-3xHA* and *UAS-Ahcy[C195S]-3xHA* flies were generated in the *attP40* landing site. Fly stocks used in this study are listed in Supp Table 1. 150 male flies were collected per sample (n = 3) for RNA-seq and CUT&RUN analysis.

### SAM and SAH Metabolite Analysis

Cells and/or heads were resuspended in methanol with internal standards (SAM-d3 and SAH-d4) and homogenized. After centrifugation, the supernatant was analyzed using an Agilent LC-MS/MS system with a Waters HILIC column separated using a pH 3 gradient, data acquired in positive electrospray ionization mode. Agilent Masshunter software was used for analysis.

### S2 Cell Transfection and Co-Immunoprecipitation

S2 cells were transiently co-transfected with actin-GAL4 and pUAST-Ahcy, AhcyL1, or AhcyL2 vectors, 3xHA or V5 tagged as described. Cells were harvested and lysed with NP-40 buffer (50 mM Tris-HCl pH 8.0, 150 mM NaCl, 1% NP-40, 10% glycerol) containing protease inhibitors and proteins immunoprecipitated using Anti-HA Magnetic beads (Thermo Scientific, Cat #88836). Western blotting was performed usinganti-HA-HRP (Roche, Cat #12013819001), anti-V5-HRP (Life Technologies, Cat #R961-25), anti-alpha-Tubulin (DSHB, Cat AA4.3).

### MM(PEG)_24_ Conjugation

Total protein from fly heads was TCA-precipitated and MM(PEG)_24_ conjugation performed as described in (Pant et al. 2021). Proteins were visualized via western blot using Anti-HA-HRP (Roche, Cat #12013819001).

### Yeast Two-hybrid Assay

Yeast two-hybrid analysis was performed with the Matchmaker Gold Yeast two-hybrid system (Clontech).

### Ahcy Label-Free Kinetic Assays

Codon optimized Ahcy was purified from *Escherichia coli* using a Nickel HiTrap HP Column (GE Healthcare) on an AKTA pure FPLC system followed by size exclusion chromatography with a Superdex S200 10/300 GL column (Cytiva). The protein was incubated with Tobacco-etch virus (TEV) protease to cleave the His tag. 60 nM Ahcy stock solutions were preincubated with 2 mM NAD+ and diluted in 100 mM pH 8.0 phosphate buffer with 1 mM EDTA and 1 mM TCEP. Reactions were started by mixing Ahcy and SAH at 37°C, with final concentrations of 30 nM Ahcy and SAH ranging from 6.25 to 400 µM. Reaction progress was monitored by aliquoting and quenching with heat. Guanosine was added as an internal standard before analysis via DESI mass spectrometry. Reaction rates were used for kinetic characterization through non-linear curve fitting in OriginPro. For H_2_O_2_ treatment, TCEP was not utilized in the assay buffer, and Ahcy stock solutions were preincubated with 10 mM H_2_O_2_ on ice and repurified before use. Kinetic parameters were plotted following a substrate inhibition model using equation y = *V*_*max*_ * x / (*K*_*m*_ + x * (1 + x / *K*_*i*_))

### Ahcy Crystallization

Frozen Ahcy protein was thawed out and incubated with TEV protease in the ratio of 1:100 (TEV: Ahcy) for overnight at 4°C. Ahcy with its cleaved His tag was then concentrated down to 1 mL and applied to a 24mL Superdex 200 10/300 GL column (Cytiva) to separate Ahcy from TEV protease and remove any protein aggregates. 12.5 mg/mL protein was added to crystallization reservoir (.1 M Bis-Tris pH 6.5 with 16% PEG MME 5000) in a 24-well hanging-drop tray. X-ray data was collected on rotating Copper anode source at Purdue Center for Macromolecular Protein Crystallography and X-ray diffraction Core. Final protein model and refinement were done using Phenix molecular replacement using human Ahcy (PDB ID: 5W49) as the basis.

### RNA Extraction and RT-qPCR

Heads were dissected from male flies, and RNA was extracted using Zymoprep Direct-zol RNA MicroPrep kit (Zymo Research, Zurich, Switzerland, Cat # R2050). cDNA was generated from 100 ng of RNA using Episcript Reverse Transcriptase (Epicentre, Petaluma, CA, USA) and random hexamer primers. Primer sequences are listed in Supp. Table 2.

### RNA-seq and Analysis

Photoreceptor nuclei were affinity enriched for RNA isolation as previously described (Jauregui-Lozano et al. 2021). A full protocol is available at: dx.doi.org/10.17504/protocols.io.buiqnudw. Briefly, the GFP^KASH^ (Klarsicht/Anc-1/Syne homology) domain, which localizes to the outer nuclear membrane, was used to immunoprecipitate tagged nuclei. Nuclear RNA was purified using ZYMO Direct-zol RNA Microprep Kits (#R2061). RNA library were prepared using NEBNext® Ultra™ II Directional RNA Library Prep Kit for Illumina (E7760S). rRNA was depleted using NEBNext® RNA Depletion Core Reagent Set with RNA Sample Purification Beads (E7870L), and customized rRNA probes. Libraries were sequenced using NovaSeq X Plus. Analysis was performed as previously described using edgeR v4.4.2 with limma (Escobedo et al. 2022).

### CUT&RUN and Analysis

CUT&RUN was performed on bead-bound nuclei as described in (Meers et al. 2019) using pAG-MNase fusion protein (Addgene plasmid #123461) purified using the Pierce cobalt kit (ThermoFisher, Pierce^TM^ His Protein Interaction Pull-Down Kit, Cat. #21227). The following antibodies were used for CUT&RUN with a 20-minute pAG-MNase activation time: anti-H3K4me3 (EpiCypher, 13-0060), anti-H3K27me3 (EpiCypher, 13-0055) and IgG (EpiCypher, 13-0042). CUT&RUN libraries were prepared using the NEBNext® Ultra^TM^ II DNA Library Prep Kit for Illumina®. Libraries were sequenced on the Illumina NovaSeq X Plus platform. Three samples (mCherryRNAi untreated H3K4me3 replicate 2, mCherryRNAi untreated H3K27me3 replicate 1, and AhcyRNAi untreated H3K4me3 replicate 2) were removed after preliminary analysis due to poor data quality. Pair-ended reads were processed similarly as RNA-seq data, except bowtie2 (Langmead and Salzberg 2012) was used for alignment with the *–dovetail* argument. Data is shown as CPM-normalized and subtracted for respective IgG control. Metaplots were generated using deepTools (Ramírez et al. 2016) based on custom GTF files represented unique used transcripts.

### Retina Staining and Analysis

Dissection and staining using Phalloidin 594 was conducted as previously described (Escobedo et al. 2022). Images were taken on a Nikon A1RSi. Rhabdomere loss was quantified blindly by scoring 20 ommatidia per sample.

## Data Availability

High-throughput sequencing data is available at Gene Expression Omnibus (GEO). GSE289458: Light-stressed D5 mCherry RNAi and Ahcy RNAi RNAseq (12 samples); GSE294368: Light-stressed D5 mCherry RNAi and Ahcy RNAi CUT&RUN (33 samples); X-ray crystal structure PDB submission in progress.

## Acknowledgements

We thank the Weake lab for their comments on the manuscript. We also thank Amber Jannasch and the Metabolite Profiling Facility at the Bindley Bioscience Center for processing the metabolite samples.

## Author Contributions

S.C.S contributed targeted metabolite studies, MM(PEG)_24_ conjugation studies, co-immunoprecipitation in S2 cells, RT-qPCR, light-stress RNA-seq, and confocal microscopy data. G.M. contributed to data analysis of RNA-seq. S.C.S and K.S. contributed to production of recombinant proteins and generating crystals for X-ray crystallography. K.S. and A.D.M contributed to analysis of crystal structure. S.C.S, N.M.M., Y.F., and R.G.C. contributed to HT-DESI-MS. N.M.M. contributed to analysis of HT-DESI-MS. M.N.M contributed to preparation and analysis of CUT&RUN of H3K4me3 and H3K27me3. C.C.K. contributed to Y2H. M.M.J contributed to co-immunoprecipitation in S2 cells. S.C.S and V.M.W. contributed to bulk data analysis, writing, and editing.

---

**Supplemental Figure 1.**
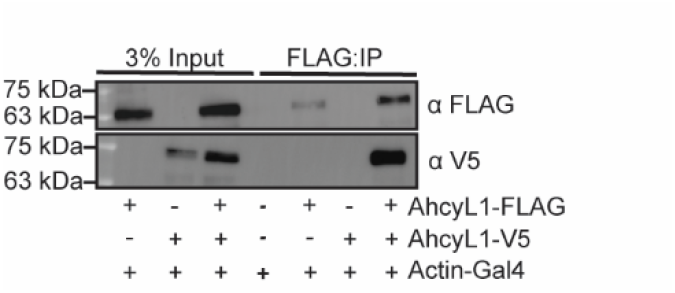
*(A)* Co-immunoprecipitation of overexpressed epitope-tagged AhcyL1 in S2 cells. Cells were co-transfected with Actin-Gal4 and AhcyL1 tagged constructs and were analyzed via western blot.

**Supplemental Figure 2.**
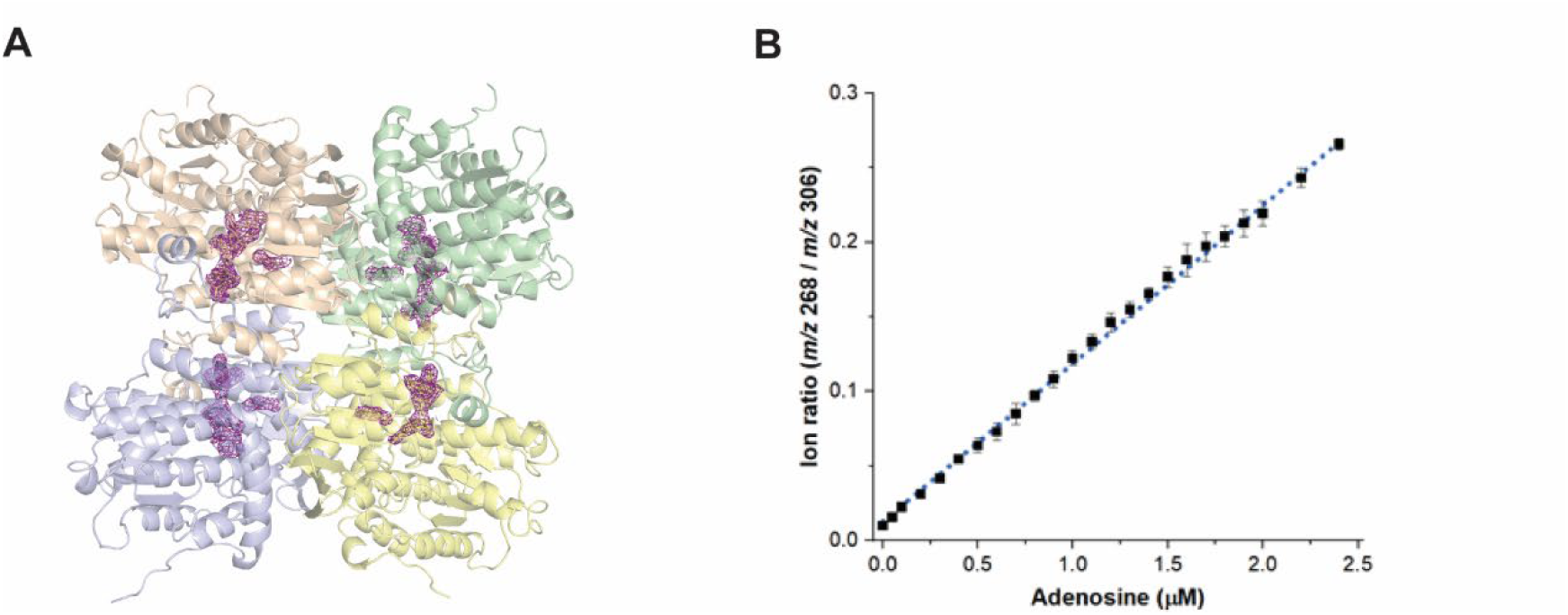
*(A)* X-ray crystal structure of WT *Drosophila* Ahcy at 2.7 Å with ligand density shown *(purple)*. Subunit chain A *(orange)*, chain B *(green)*, chain C *(blue)*, and chain D *(yellow). (B)* Calibration curve for the determination of adenosine (m/z 268) using guanosine (m/z 306) as internal standard via HT-DESI-MS. [M+H]+ and [M+Na]+ ion species were monitored for adenosine and guanosine, respectively. Six independent replicates, each with eight technical replicates, were analyzed for every calibration solution (>1,000 samples). The linear fit obtained (R2 = 0.995) is also included. Data points indicate averages and error bars standard deviations (n = 48). Note the high precision (all coefficients of variance, CVs, below 12%) and sensitivity (limit of detection, LOD, 120 nM) of the methodology.

**Supplemental Figure 3.**
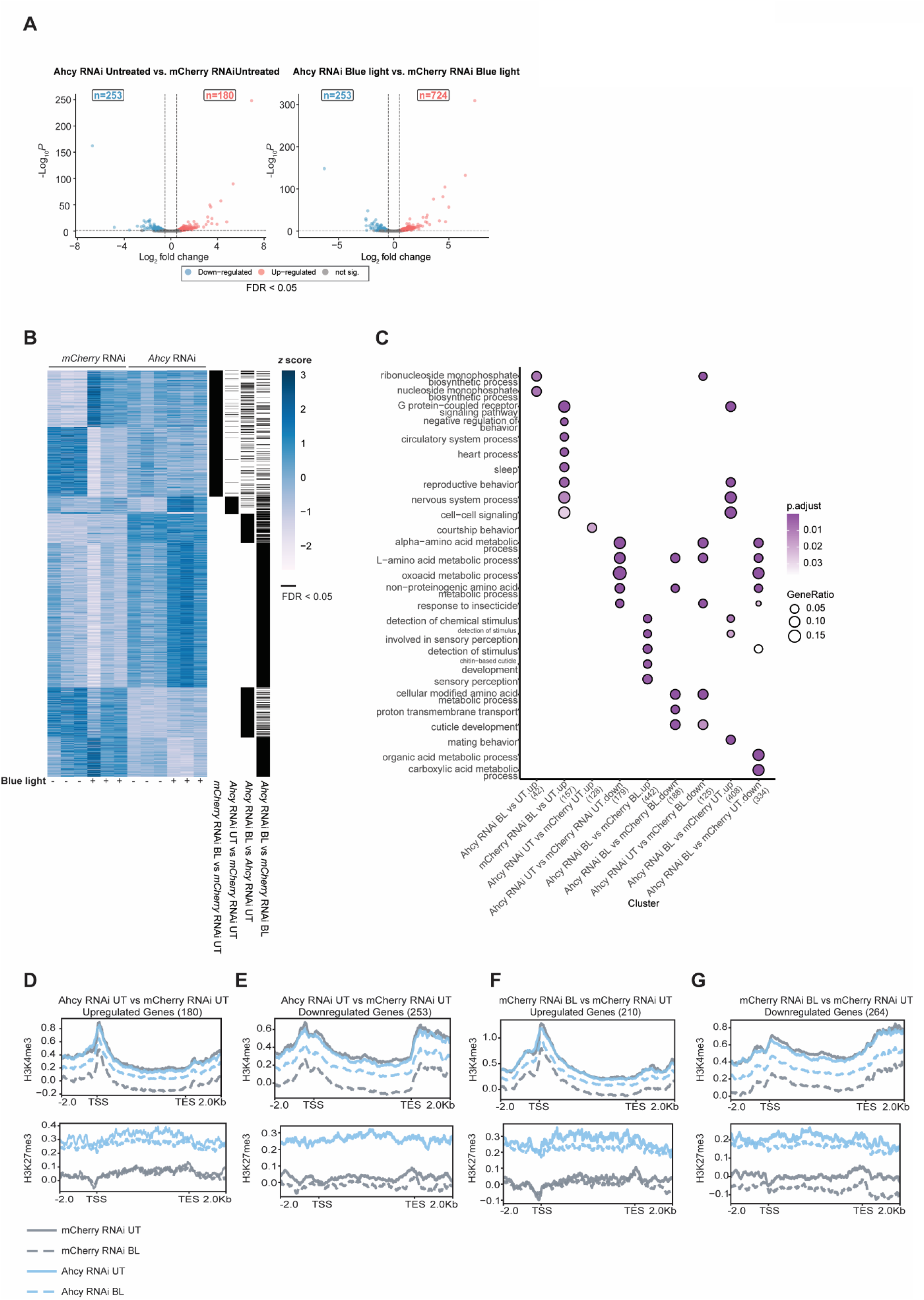
*(A)* Volcano plots displaying significantly up- and down-regulated genes in either *mCherry* RNAi flies or *Ahcy* RNAi flies exposed to blue light relative or untreated controls. FDR < 0.05, log_2_(fold-change) ≥ 1 or ≤ −1). *(B)* Heatmap showing genes that are differentially expressed genes (DEGs) from *mCherry* RNAi flies exposed to blue light relative to untreated controls, *Ahcy* RNAi flies exposed to blue light relative to untreated controls, *Ahcy* RNAi untreated flies relative to *mCherry* RNAi untreated flies, and *Ahcy* RNAi flies exposed to blue light relative to *mCherry* RNAi flies exposed to blue light. Colors indicate *z* score of relative expression in biological replicates for each genotype and treatment. *(C)* Dot plots of all enriched Gene Ontology (GO) terms. *(D-G)* Gene metaplots representing mean normalized counts for CUT&RUN of indicated histone methyl marks for up- and downregulated genes (n = 3).

